# *D*_GEN_: A Test Statistic for Detection of General Introgression Scenarios

**DOI:** 10.1101/348649

**Authors:** Ryan A. Leo Elworth, Chabrielle Allen, Travis Benedict, Peter Dulworth, Luay Nakhleh

## Abstract

When two species hybridize, one outcome is the integration of genetic material from one species into the genome of the other, a process known as introgression. Detecting introgression in genomic data is a very important question in evolutionary biology. However, given that hybridization occurs between closely related species, a complicating factor for introgression detection is the presence of incomplete lineage sorting, or ILS. The *D*-statistic, famously referred to as the “ABBA-BABA” test, was proposed for introgression detection in the presence of ILS in data sets that consist of four genomes. More recently, *D*_FOIL_—a set of statistics—was introduced to extend the *D*-statistic to data sets of five genomes.

The major contribution of this paper is demonstrating that the invariants underlying both the *D*-statistic and *D*_FOIL_ can be derived automatically from the probability mass functions of gene tree topologies under the null species tree model and alternative phylogenetic network model. Computational requirements aside, this automatic derivation provides a way to generalize these statistics to data sets of any size and with any scenarios of introgression. We demonstrate the accuracy of the general statistic, which we call *D*_GEN_, on simulated data sets with varying rates of introgression, and apply it to an empirical data set of mosquito genomes.

We have implemented *D*_GEN_ and made it available, both as a graphical user interface tool and as a command-line tool, as part of the freely available, open-source software package ALPHA (https://github.com/chilleo/ALPHA).

## 1 Introduction

Hybridization—the interbreeding of individuals from two “different” species, or populations—has been recognized as an important evolutionary process underlying genomic diversification and species adaptation [1, 2, 22, 14, 15, 21, 8, 19, 16, 26]. Immediately upon interbreeding, each chromosome in the hybrid individual has a single source—the genome of one of the two parents. However, after multiple generations of backcrossing and recombination, the genomes of descendants of the hybrid individual turn into mosaics of genomic segments, each having a genealogy that could potentially differ from that of other segments (Fig. 1). The integration of genetic material from two different species into the genome of an individual is called *introgression*.

**Figure 1:**
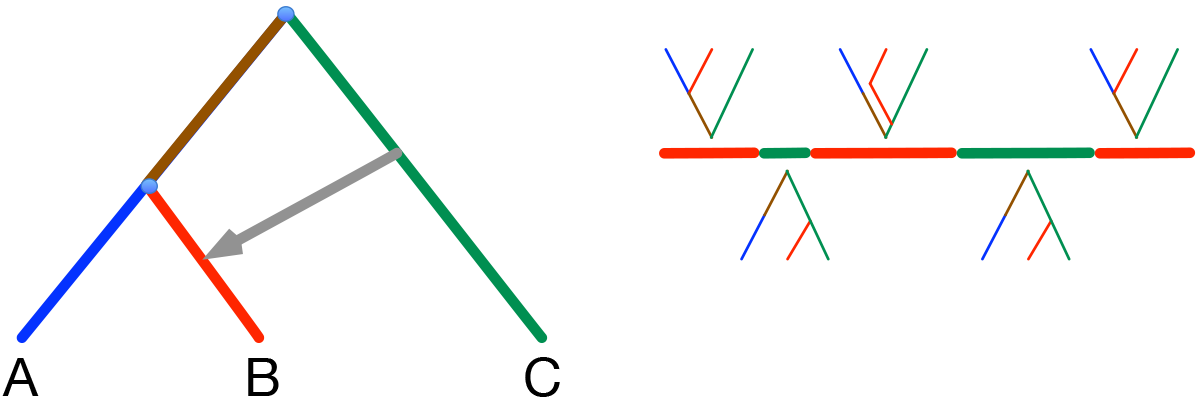
Hybridization and introgression. Left) A phylogenetic network modeling the evolutionary history of three species (or, populations) A, B, and C. Species C split from the most recent common ancestor of A and B, and hybridization between (an ancestor of) C and (an ancestor of) B occurred. (Right) Due to hybridization and backcrossing, the genome of an individual in species B is a mosaic with different genomic segments having different genealogies. In particular, the genealogy of the middle segment involves incomplete lineage sorting.

The discordance among the genealogies of different genomic segments could be used as a signal to detect introgression. For example, in the case of Fig. 1, the presence of some genealogies that place B closer to A than to C and others that place B closer to C than to A could indicate a potential hybridization event between B and C. However, a complicating factor in introgression detection is that incomplete lineage sorting, or ILS, could also be at play in cases where hybridization has occurred. ILS occurs when lineages from related populations fail to coalesce within the ancestral population, giving rise to the possibility that some lineages coalesce with others from farther populations. Mathematically, this process is often modeled by the multispecies coalescent [10, 24, 17, 5].

One class of methods for detecting hybridization and introgression, including in the presence of ILS, is to infer phylogenetic networks from the data of multiple unlinked loci sampled across the genomes. Indeed, several methods were introduced recently for this task [31, 29, 27, 23, 25, 32, 34, 33]. While providing accurate results, these methods are computationally very demanding.

A different approach is to use the so called *D*-statistic [9, 6], which infers the presence of introgression based on significant deviation from equality between the frequencies of two site patterns in a 3-taxon (plus an outgroup) data set (details below). More recently, Pease and Hahn [18] introduced *D*_FOIL_, which extends the *D*-statistic to detect introgression in a 5-taxon scenario (4 taxa plus an outgroup). The extension from three to four taxa involved a detailed analysis of site patterns and resulted in a set of statistics that, when combined, would aid in the detection of introgression. However, as stated, that work of Pease and Hahn extended the *D*-statistic from three to four taxa. Both the *D*-statistic and *D*_FOIL_ are examples of the use of phylogenetic invariants to detect deviation from the expected frequencies of site patterns under a neutral coalescent model with no gene flow. Similarly, the HyDe software package [3] implements an invariants-based method for identifying hybridization [13].

A major question is: Can one devise a statistic that is general enough to apply (the computational complexity issue aside) to data sets with any number of genomes and any set of postulated hybridization events?

In this paper, we address this question by showing that the phylogenetic invariants underlying both the *D*-statistic and *D*_FOIL_ could be generated automatically by contrasting gene tree distributions under the null multispecies coalescent [5] and the alternative multispecies network coalescent [30, 31]. Based on this observation, we devise an algorithm that automatically generates a statistic for detecting introgression in any evolutionary scenario. It is important to note, though, that as the number of genomes and number of postulated hybridization events increase, computing the statistic becomes computationally very demanding.

Our method, which we call *D*_GEN_, is implemented in the publicly available, open-source software package ALPHA [7]. We demonstrate the accuracy of the method on simulated data sets, as well as its applicability to an empirical data set of mosquito genomes.

## 2 Methods

### 2.1 The *D*-statistic

Consider the species tree (((P1,P2),P3),O) in Fig. 2, which shows the evolutionary history of three species, or populations, P1, P2, and P3, along with an outgroup O. The significance of an outgroup in this scenario is that for any genomic site, the state that the outgroup has for that site is assumed to be the ancestral state of all three species P1, P2, and P3. We denote by A the ancestral state and by B the derived state.

Assuming all lineages from P1, P2, and P3 coalesce before any of them could coalesce with a lineage from O, there are three possible gene trees topologies, which are shown inside the branches of the species tree in Fig. 2, and are given by (((P1,P2),P3),O), (((P1,P3),P2),O), and (((P2,P3),P1),O). The probabilities of these three gene tree topologies when gene flow is excluded (but incomplete lineage sorting is accounted for) are, respectively, 1 − (2/3)*e*^−*t*^, (1/3)*e*^−*t*^, and (1/3)*e*^−*t*^, where *t* is the length, in coalescent units, of the branch that separates the splitting of P3 from the ancestor of P1 and P2 [4]. Clearly, the latter two gene tree topologies (those that are discordant with the species tree) have equal probabilities. Taking the two patterns BABA and ABBA to correspond to gene trees (((P1,P3),P2),O) and (((P2,P3),P1),O), then their expected frequencies in the absence of gene flow are equal.

However, when gene flow from P3 to P2 occurs and is modeled as an instantaneous event with probability *γ* (*γ* here is taken to represent the fraction of genomes in P2 that originated from P3 through gene flow), then the probabilities of the three gene tree topologies (((P1,P2),P3),O), (((P1,P3),P2),O), and (((P2,P3),P1),O) become, as derived in [28], 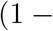 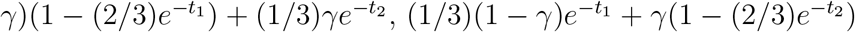, and 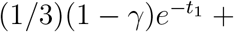 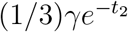, respectively, where *t*_1_ and *t*_2_ are the branch lengths, in coalescent units, in the phylogenetic network of Fig. 2. Now, with gene flow accounted for, when *γ* ≠ 0(and *t*_2_ > 0), the expected frequencies of the two patterns BABA and ABBA are no longer equal. Thus, denoting by *N_X_* the number of times site pattern *X* appears in a genomic data set, the *D*-statistic was defined as [6]

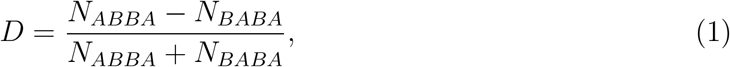

and the significance of the deviation of *D* from 0 is assessed. Under no gene flow, we expect *D* ≈ 0 (we do not write *D* = 0 since the counts in eq. (1) are estimated from actual data and might not match the theoretical expectations exactly), and in the presence of gene flow, we expect *D* to deviate significantly from 0. Furthermore, when *D* > 0, it indicates introgression between P2 and P3 (in either or both directions), and when *D* < 0, it indicates introgression between P1 and P3.

**Figure 2:**
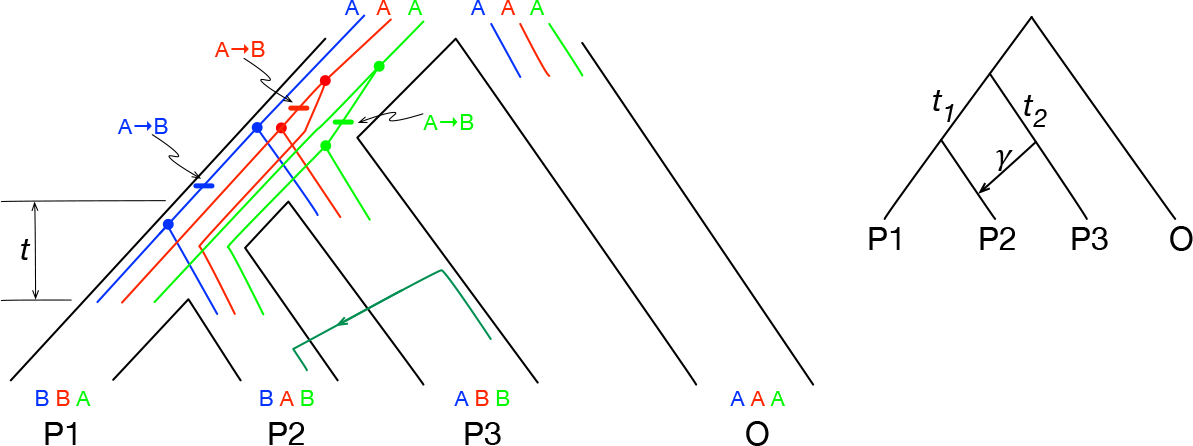
Illustration of the*D*-statistic. (Left) The demographic structure of three populations, P1, P2, and P3, along with an outgroup O, is shown. Patterns of the three parsimony-informative mutations (A→B) are shown for biallelic sites with states A and B, each mapped onto a different genealogy. The three genealogies give rise to patterns BBAA, BABA, and ABBA for the four taxa, respectively, when the taxa are listed in the order P1-P2-P3-O. The dark green arrow indicated gene flow from P3 to P2, which would result in excess of pattern ABBA. (Right) The phylogenetic network modeling the evolutionary history of the populations in the presence of gene flow from P3 to P2, where the gene flow is modeled as an instantaneous unidirectional event.

To extend the *D*-statistic from the scenario depicted in Fig. 2 to the case of five taxa (four populations and an outgroup), Pease and Hahn [18] identified sets of site patterns that are expected to have equal frequencies under a no gene flow scenario but different frequencies when gene flow occurs. Next we show how to derive a general *D*-statistic that applies to a species phylogeny and any set of gene flow events, thus overcoming the need to derive a specialized *D*-statistic for individual evolutionary histories.

### 2.2 Towards the General Case

Let *X* be a set of taxa *X*_1_, *X*_2_, …, *X_n_*, where *X_n_* is assumed to be an outgroup whose state A for a given bi-allelic marker is assumed to be the ancestral state. Then, for a given marker, a site pattern *s* is a sequence of length *n* where *s_i_* (1 ≤ *i* < *n*), the state of the site in the genome of *X_i_*, is either A or B.

Let 𝒢 be the set of all rooted, binary gene trees on the *n* taxa *X*_1_, …, *X_n_*. For a site pattern *s*, there might be multiple trees in 𝒢 that are *compatible* with *s*; that is, trees on which the pattern *s* could have arisen in the presence of a single mutation (the infinite-sites assumption). We denote by 𝒢(*s*) the set of all trees in 𝒢 that are compatible with pattern *s*. While the size of 𝒢 only depends on the number of taxa *n*, the size of 𝒢(*s*) for a given *s* also depends on the number of ancestral versus derived alleles represented in *s*. For a given *s* with *n* total taxa and *β* taxa having the derived state (a ‘B’ instead of a ‘A’ in the site pattern), the size of 𝒢(*s*) will be the number of rooted, binary trees on *β* taxa times the number of rooted, binary trees on *n* − *β* + 1 taxa. Given a species phylogeny Ψ, we have

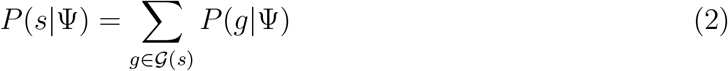

where *P* (*g*|Ψ) is the probability mass function (pmf) of [4] when Ψ is a species tree and the pmf of [28] when Ψ is a phylogenetic network.

Assuming independence among sites, the expected number of occurrences of site pattern *s* in a genomic data set given species phylogeny Ψ, denoted by 𝔼(*n*(*s*)), is given by *n·P* (*s*|Ψ). Using this notation, a general statistic for detecting introgression proceeds as follows. Let Ψ be the species tree that corresponds to the evolutionary scenario of no gene flow. Let Ψ*’* be the phylogenetic network that is obtained by adding to Ψ the gene flow events (instantaneous events represented by horizontal edges) to be tested on Ψ. For example, the phylogenetic network in Fig. 2 is obtained by adding the gene flow event from P3 to P2. A general *D*-statistic, *D*_GEN_, is then computed as follows:

1. Let *S* be the set of all distinct parsimony-informative site patterns.
2. Parameterize Ψ and Ψ*’* so that they define probability distributions on gene tree topologies.
3. For every site pattern *s* ∈ *S*, compute *P* (*s*|Ψ) and *P* (*s|*Ψ*’*).
4. Let 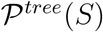 be the partition of set *S* induced by the equivalence relations {(*s*_1_, *s*_2_): *P* (*s*_1_|Ψ) = *P* (*s*_2_|Ψ}.
5. Let 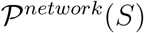 be the partition of set *S* induced by the equivalence relations (*s*_1_, *s*_2_): *P* (*s*_1_|Ψ*’*) = *P* (*s*_2_|Ψ*’*)}
6. Let 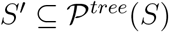 where *Y* ∈ *S’* if and only if *Y* ⊈ *Z* for any 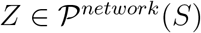. In other words, *Y* is an element of *S’* if it consists of a set of site patterns that all have equal probabilities under Ψ but not equal probabilities under Ψ*’*.
7. Let 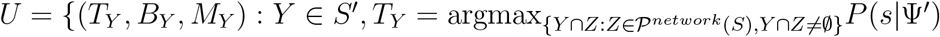 and 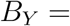 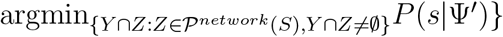 where *s* is an arbitrary element of *Y* ∩ *Z*, and *M*_*Y*_ = *Y* − (*T*_*Y*_ ∪ *B*_*Y*_). Put simply, site pattern probabilities that were previously equal in the tree case become totally ordered in the network case and can be divided into sets based on their new relation to one another. In other words, as an equivalence class *Y* ∈ *S’* is refined by the elements of 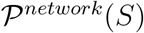, *T*_*Y*_ and *B*_*Y*_ are the two subsets of site patterns in *Y* with the highest and lowest probabilities, respectively, and *M*_*Y*_ is the set of remaining site patterns.
8. *D*_GEN_ = (Σ_(*T,B,M*)∈*U*_ *N*_*T*_ − *N*_*B*_) / (Σ_(*T,B,M*)∈*U*_ *N*_*T*_ + *N*_*B*_ + 2*N*_*M*_) where, as above, *N*_*T*_ is the number of times site patterns in *T* appear in the genomic data set (and similarly for *N*_*B*_ and *N*_*M*_).
9. Similar to [18], calculate the *χ*^2^ goodness of fit (df=1) using

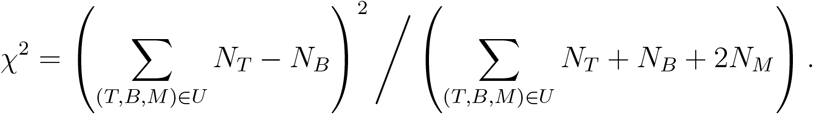

Applying this algorithm to the case illustrated in Fig. 2, we have

- 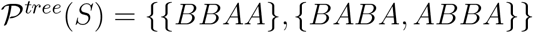.
- 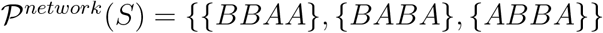.
- Step (6) returns *S’* = {{*BABA*, *ABBA*}}.
- Step (7) returns *U* = {({*BABA*},{*ABBA*})}which, indeed, is the *D*-statistic in the case of three taxa.

In Step (2) of the algorithm, we parameterize Ψ (and Ψ*’*) by trying branch lengths (in coalescent units) in the set of values 0.5, 1.0, 2.0, 4.0 and the set *S’* (in Step (6)) is determined based on the sets of site patterns whose equality does not break across the different settings of branch lengths. For the inheritance probability, we set it to 0.9.

In Step (7), if for an element (*T*, *B*, *M*) of *U*, we have |*T*| ≠ |*B*|, we remove (arbitrarily) elements from the larger of the two sets to make them of equal size. Here, |*T*| is distinguished from *N*_*T*_ as |*T*| represents how many site patterns are contained in set T (the cardinality of set T), whereas *N*_*T*_ represents the occurrence of the site patterns in T in an actual multiple sequence alignment.

For the *χ*^2^ test, we used a threshold of 0.01 on the *p*-value to determine significance. That is, if the *p*-value is smaller than 0.01 we considered support for introgression to be statistically significant; otherwise, it is not. Given that our formulation bases its determination of whether introgression is present or not off of significant deviations of *D*_GEN_ away from zero, sign changes are treated equivalently and thus one could equivalently choose to take the absolute value of *D*_GEN_. In our implementation of *D*_GEN_ we have chosen to leave the sign. More information on this is given in the discussion section.

**Why not contrast the site pattern distribution to the known distribution of gene trees?** One question that might arise is: Why do we not use a *χ*^2^ test to compare the two distributions—the empirical one and the theoretical one; that is,

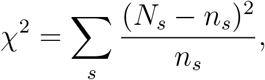

where the sum is taken over all distinct site patterns *s*, *N*_*s*_ is the observed count of site pattern *s*, and *n*_*s*_ = *n* · *P* (𝒢(*s*) | Ψ). The problem with this approach is that to compute *n*_*s*_, we need knowledge of the parameters (branch lengths) of the species phylogeny, which are unknown in this case. One potential remedy to this limitation is to first estimate the species tree parameters from the data, say under maximum likelihood, and then use this parameterized model to compute the *n*_*s*_ frequencies. However, it is unknown how the estimated parameters compare to the (unknown) true values when gene flow had occurred but the assumed topology in the estimation is a tree. In our solution above, this problem is remedied by not focusing on the parameter values in an absolute sense, but rather use arbitrary settings to find the site patterns whose relative frequencies change between a model of no gene flow and another with gene flow.

## 3 Results

### 3.1 Simulations

We first studied the performance of our method on the five-taxon scenario studied in [18] and given by species tree Ψ_1_ in Fig. 3. All simulations share the same values for several parameters. As in [18], we have a constant fixed population size of *N*_*e*_ = 10^6^ and recombination rate of *r* = 10^−8^. We also use a fixed mutation rate of *μ* = 7 × 10^−9^. In our simulation pipeline, we first generate gene trees for a 50kbp multiple sequence alignment using **`ms`** [11], followed by simulating the sequences under the Jukes-Cantor model of evolution [12] using **`seq-gen`** [20]. In other words, the sequences are evolved under a finite-sites model. The parameter values were chosen primarily to accomplish the two goals of being similar to relevant past work as well as being biologically relevant. An example of the full commands for this pipeline, before adding any reticulations is as follows:

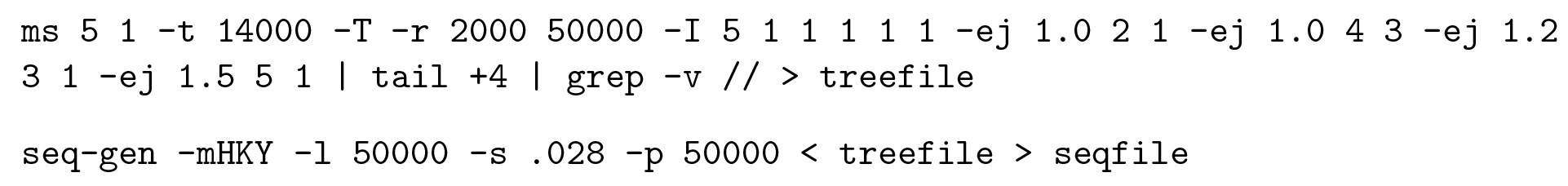

We then added a migration event between P1 and P3 at time 0.5, with varying migration rates, and calculated our *D*_GEN_ statistic on the resulting genomic data sets; results are shown in Fig. 3. As the results show, *D*_GEN_ performs very well at determining the presence of introgression in data sets. In particular, when the data evolved with no migration (migration rate 0), the *D*_GEN_ values hardly deviate from 0, and when the migration rate is non-zero, the method detects the presence of introgression with a strong deviation from 0. These results are consistent with the performance of *D*_FOIL_ [18].

**Figure 3:**
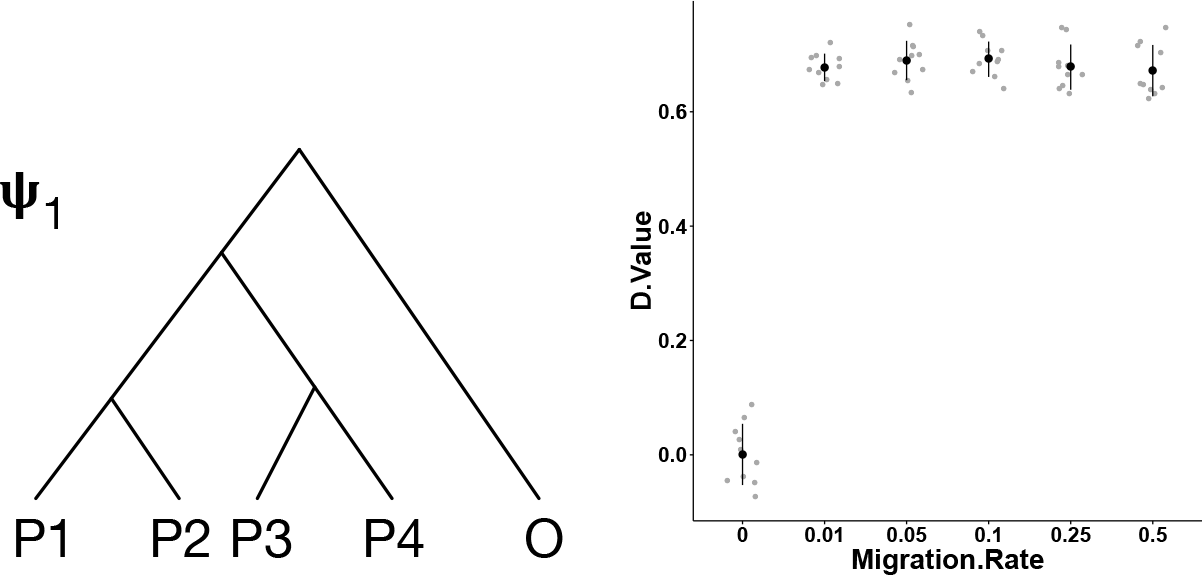
5-taxon simulation results. (Left) A 5-taxon species tree. The two most recent divergence events are set at 1.0 coalescent units, the divergence time of the ancestor of all in-group taxa (P1–P4) is set to 1.2 coalescent units, and the time of the root node is set to 1.5 coalescent units. A migration event between P1 and P3 at time 0.5 was added to species tree. (Right) Values of *D*_GEN_ on data sets with varying migration rates. Each point corresponds to a *D*_GEN_ value whose *p*-value was lower than 0.01 obtained from a different data set simulated under the same settings. The dark dots correspond to the mean and the lines correspond to 1 standard deviation around the mean.

Next, we considered cases beyond that of five taxa (i.e., cases not possible with either the *D*-statistic or *D*_FOIL_). We conducted simulations that show the effect of migration rate and time of the migration event on the performance of *D*_GEN_, as shown in Fig. 4.

As the results show, the *D*_GEN_ statistic performs very well at detecting introgression in this case as well. In particular, as the migration rate increases, so does the accuracy of the method. For a migration rate of 10^−6^ or higher, the method detects, with high significance,the presence of introgression. In the cases of extremely low migration rates (10^−7^ and 10^−8^),the method tends to indicate slight deviation from a no-introgression scenario.

As for varying the time of the migration event, the accuracy of the method is what one would expect. As the time between the migration event and the divergence event increases (the time of the migration event decreases), the power to detect introgression is much higher. That power starts decreasing as the migration event becomes more ancient and, as a result, less signal is present for its detection.

#### 3.1.1 Multiple Reticulations

The question we set out to investigate next is: Given that the *D*-statistic is designed to work under the assumption of a single gene flow event, how does it perform when there is more than one event? Fig. 5 shows a typical scenario for the *D*-Statistic, Scenario S1, in which the standard 4-taxon backbone tree has a single reticulation from P3 to P2. The followingfour scenarios add an extra reticulation with a high migration rate. The effect of adding these reticulations on the value of the *D*-Statistic are shown in Fig. 6.

**Figure 4:**
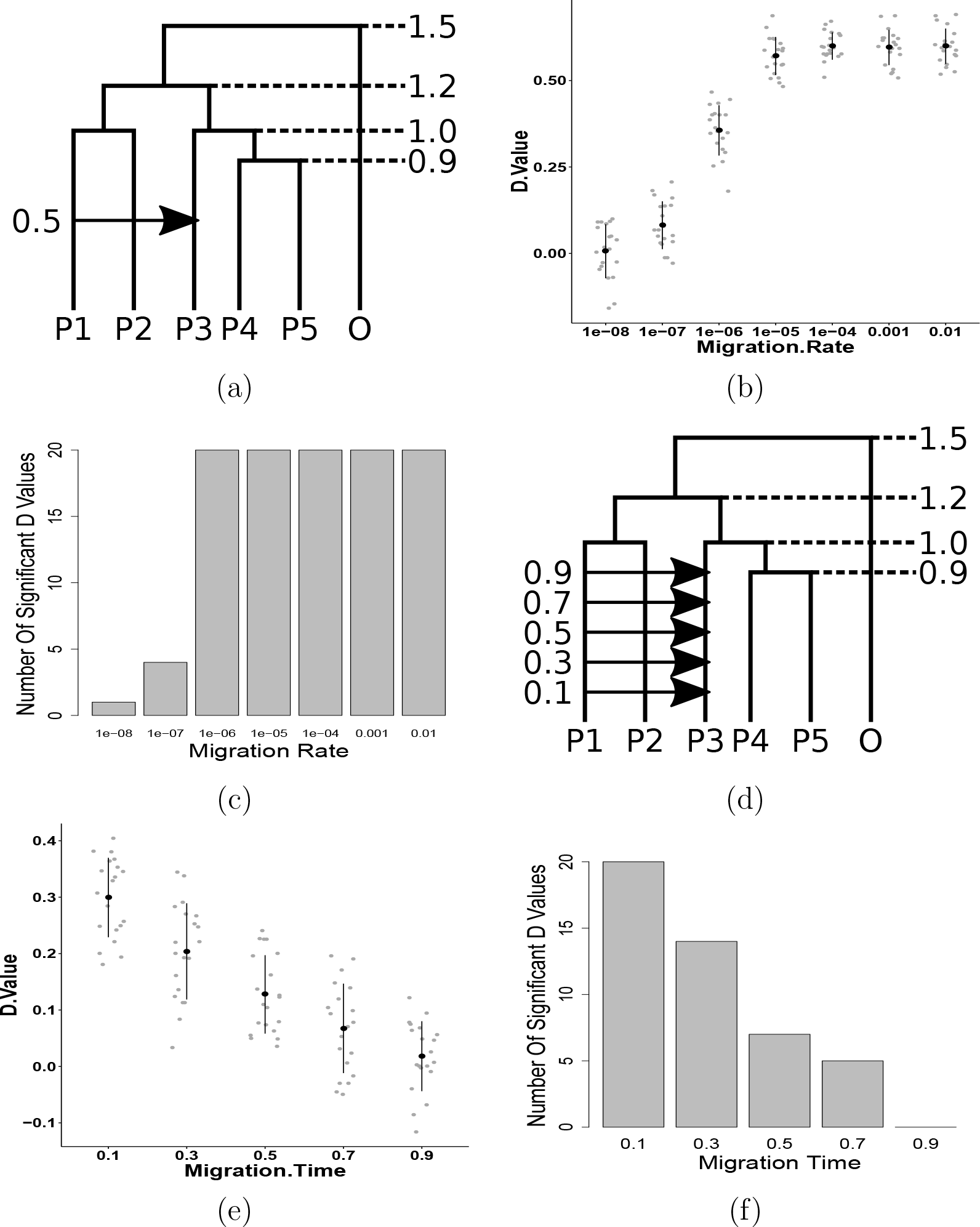
Simulation results on 6-taxon scenarios. (a) The network used for analyzing the effects of varying migration rate of a reticulation. The results of the corresponding *DG* values and the number of data sets where the *D*_GEN_ values were significant (*p*-value smaller than 0.01) are shown in (b) and (c), respectively. (d) The network used for analyzing the effects of varying the time of the migration event. The results of the corresponding *DG* values and the number of data sets where the *D*_GEN_ values were significant (*p*-value smaller than 0.01) are shown in (e) and (f), respectively.

**Figure 5:**
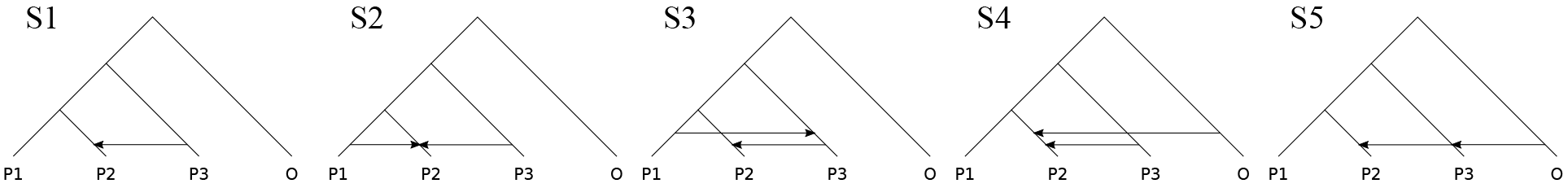
The standard D-Statistic scenario followed by four scenarios where an additional reticulation is added. S1 adds the first reticulation which is held constant throughout all scenarios and has M=0.1. The added reticulations in S2 through S5 have M=0.5.

**Figure 6:**
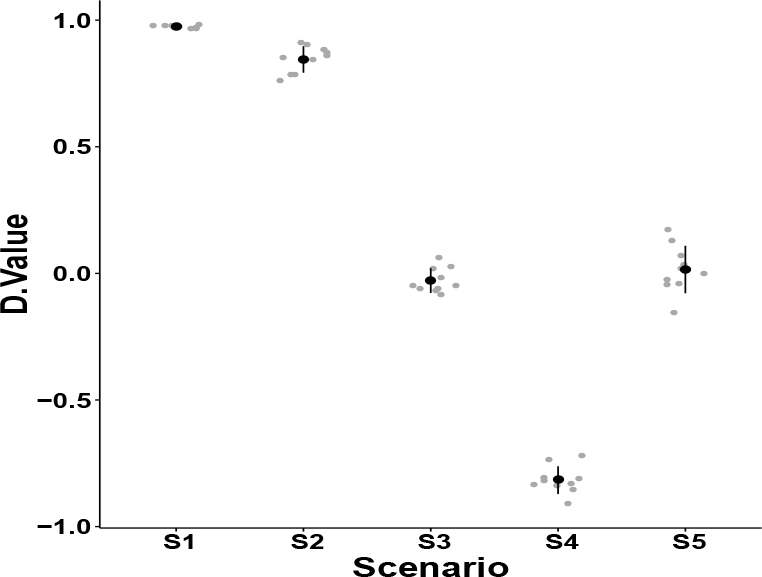
*D*-Statistic values for scenarios with a “hidden” reticulation. The scenarios S1–S5 are shown in Fig. 5.

As expected, the S1 case yields the best results, followed by the S2 case with a weaker *D* value. All other statistic values demonstrate that even in the presence of a significant migration with introgression from P3 to P2, multiple introgressions can cause that information to be lost from inference. These results show that it is important to account for multiple reticulations simultaneously, which our *D*_GEN_ statistic allows for given that by its design it is not restricted to any specific number of reticulations.

#### 3.1.2 D-Statistic Subsetting

When data sets with more than four taxa are to be analyzed by the *D*-statistic, a workaround is to subset the set of taxa into groups of four genomes (one outgroup and three in-group taxa) and conduct *D*-statistic analyses on each subset independently. Our method, being general, allows for analyzing the data set without any subsetting. The question we set out to investigate here is: Does subsetting and running the *D*-statistic on individual 4-taxon subsets equate to running *D*_GEN_ on the full data set? To answer this question, we considered the evolutionary scenario of Fig. 7.

**Figure 7:**
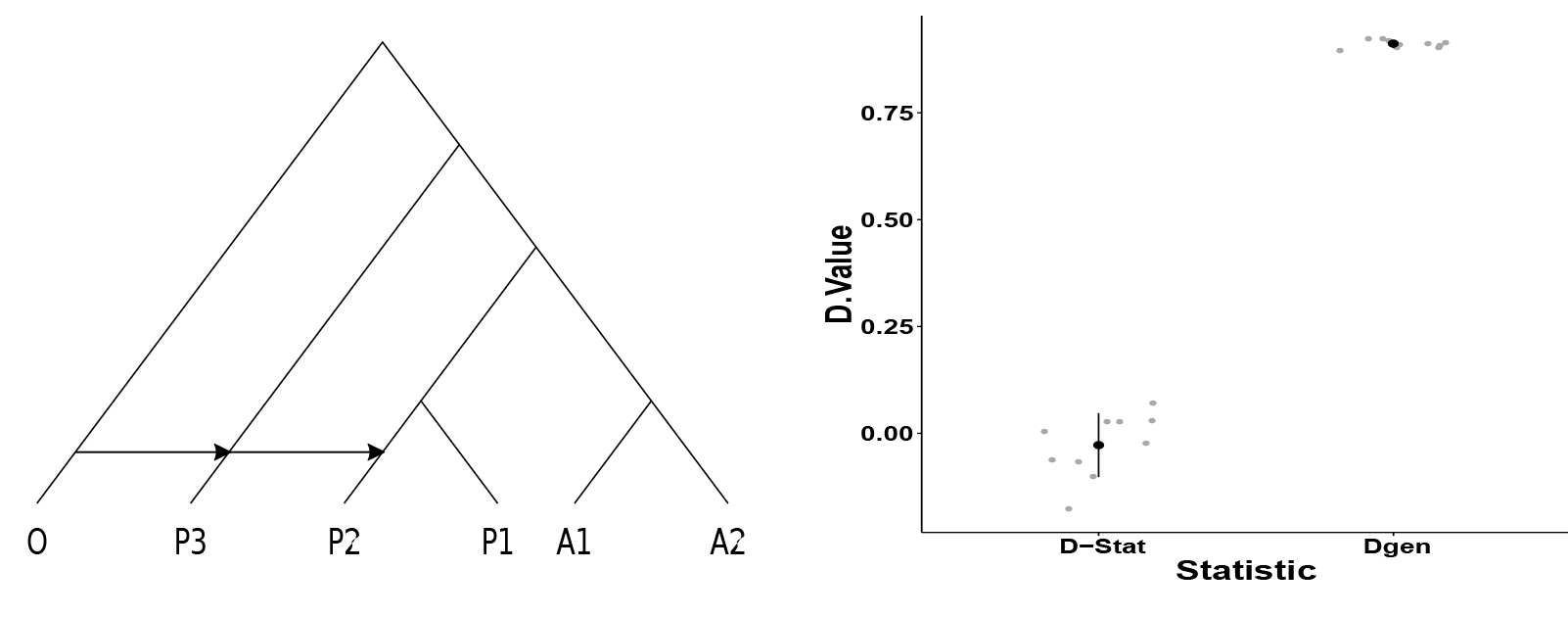
The effect of subsetting on the detectability of introgression. (Left) A 6-taxon evolutionary history with two migration events. (Right) The values of the *D*-statistic on subsets of four taxa, and *D*_GEN_ on the full data set.

In the case of the *D*-statistic, only two out of the ten simulations recovered a significant non-zero *D* value, whereas *D*_GEN_ inferred significant *D* values for all runs. This further demonstrates the need for and significance of a method that works directly on a full data set and accounting for multiple migration events.

### 3.2 Analysis Of a Mosquito Genomic Data Set

Finally, we present results from a real biological data set with six taxa. In both [8] and [26], the evolutionary history of the *Anopheles gambiae* species complex was found to be reticulate. Both studies found particularly strong signals of introgression in the 3L chromosome in an area known as the 3La inversion. This reticulation between *Anopheles quadriannulatus* (Q) and *Anopheles merus* (R) is shown in the network of Fig. 8(a) with the other species of *An. coluzzii* (C), *An. arabiensis* (A), *An. melas* (L), and *An. christyi* (O).

The results from Fig. 8(b) show that *D*_GEN_ does recover the introgressed region around the 3La inversion in comparison to the rest of the 3L chromosome, consistent with previous studies. Fig. 8(b) is an example of a figure that can be generated directly through the graphical user interface of the ALPHA toolkit [7] and is presented as generated directly from ALPHA. The figures output by ALPHA can be run on the full genome or on variable sized windows with variable sized offsets between windows. Here the window size used was 500kbp with a 100kbp offset between windows. The software can also vary the significance cutoff with which to display values as significant (green) or not significant (red). Here a significance cutoff value of 0.01 was used, as is used throughout the paper.

## 4 Discussion and Conclusions

In this paper, we extended the popular *D*-statistic to general cases of evolutionary histories of any number of taxa and any number and placement of migration events. What enabled this extension is the observation that the “ABBA-BABA” phylogenetic invariant underlying the *D*-statistic can be derived automatically by making use of the probability mass function of gene tree topologies under the multispecies coalescent and multispecies network coalescent models.

**Figure 8:**
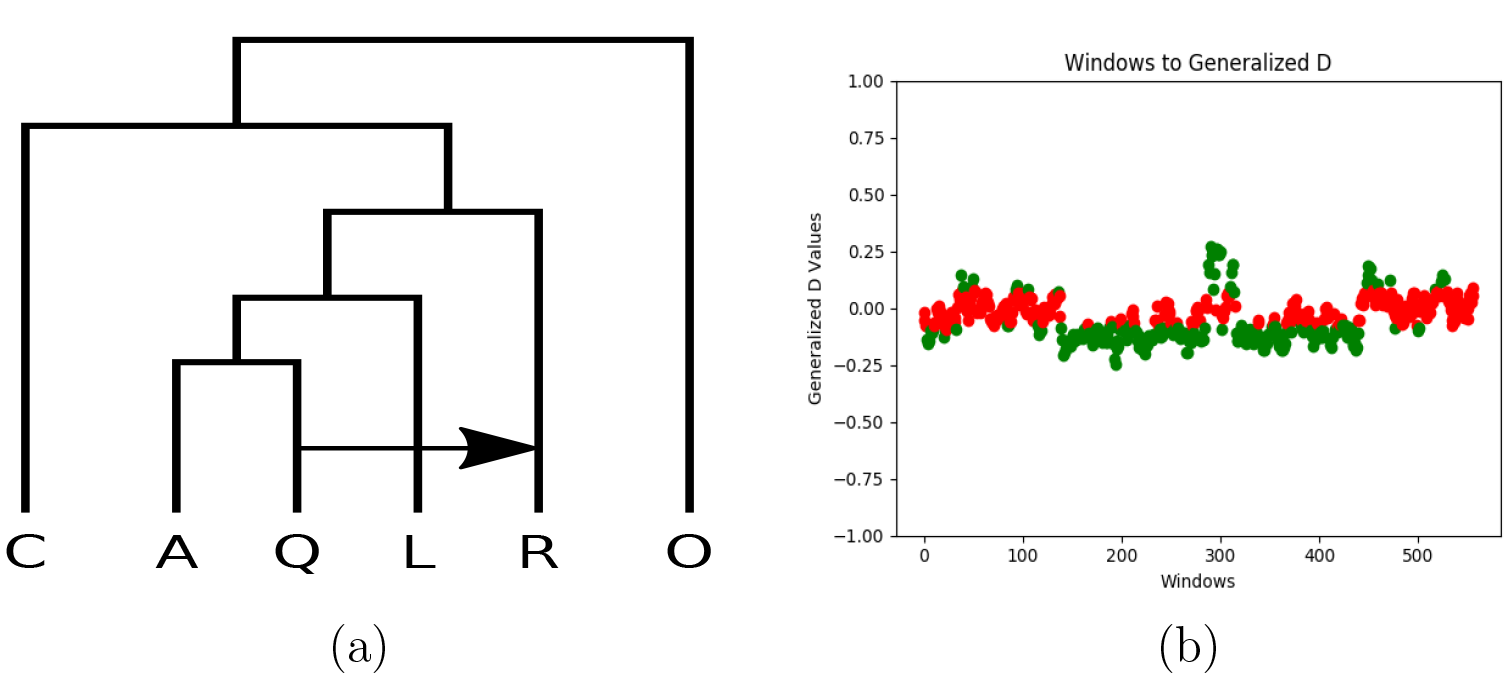
*D*_GEN_values for the 3L mosquito chromosome. (a) Evolutionary history of six mosquito genomes. (b) *D*_GEN_ analysis of the 3L chromosome with window length set to 500kbp and with a 100kbp offset between windows.

Our simulation results show that the new statistic *D*_GEN_ and method for deriving and computing it are very powerful for detecting introgression in various settings. In particular, we demonstrated that hidden migration events could negatively affect the performance of the *D*-statistic, which operates under the assumption of a single migration event. Furthermore, subsetting a data set of more than four taxa into data sets with four taxa is problematic. Our *D*_GEN_ statistic addresses these two issues by enabling the analysis of data sets with more than four taxa and more than a single migration event. While analyses in the style of the D-statistic make major assumptions, such as assuming the infinite sites model as well as ignoring dependence between sites, they are resilient to violations in these assumptions. Our results further support this given that our simulations violate both of these assumptions, having been performed under the full coalescent with recombination model with a mutation model allowing for recurrent mutation.

It is important to note that the *D*-statistic provides values that could be positive or negative. The sign of these values give an indication on the directionality of the migration in the case of four taxa. However, in the case of larger data sets, the sign of the *D*_GEN_ values is not easily interpretable in terms of directionality. It is also important to note that the actual *D* value is not the quantity of interest; rather, it is the statistical significance of its deviation from 0. There is of course, however, a strong correlation between the two.

As stated above, the method has been implemented in the ALPHA toolkit, which allows for conducting *D*_GEN_ analyses on the command-line as well as through a graphical user interface.

Finally, while we present the *D*_GEN_ statistic and its computation as a way of analyzing introgression under general evolutionary scenarios, computational complexity will become prohibitive for increasingly large, complex data sets. In particular, Step (3) in the algorithm above for computing *D*_GEN_ entails computing the probabilities of all gene tree topologies under a species tree and a phylogenetic network model. This calculation is very demanding, especially in the case of the phylogenetic network. For example, while generating *D*_GEN_ for four or five taxa takes approximately ten and forty seconds, respectively, generating it for six taxa takes thirteen minutes and for seven taxa thirty-eight hours. Fortunately, our implementation allows a *D*_GEN_ statistic to only ever need to be generated once for a particular evolutionary scenario, as the statistic itself is saved to a file that can be used on all current and future data sets for that scenario. This process of running a previously generated statistic on a new data set is, of course, computationally trivial. It will be important future work, however, to address the computational limits of *D*_GEN_ when going to arbitrarily large numbers of taxa.

